# Limited evidence for selection at the *FADS* locus in Native American populations

**DOI:** 10.1101/2019.12.11.873356

**Authors:** Iain Mathieson

## Abstract

The *FADS* locus contains the genes *FADS1* and *FADS2* that encode enzymes involved in the synthesis of long-chain polyunsaturated fatty acids (LC-PUFA). This locus appears to have been a repeated target of selection in human evolution, likely because dietary input of LC-PUFA varied over time depending on environment and subsistence strategy. Several recent studies have identified selection at the *FADS* locus in Native American populations, interpreted as evidence for adaptation during or subsequent to the passage through Beringia. Here, we show that these signals of selection are confounded by the presence of parallel adaptation–postdating their split from Native Americans–in the European and East Asian populations used in the population branch statistic (PBS) test. This is supported by direct evidence from ancient DNA that one of the putatively selected haplotypes was already common in Northern Eurasia at the time of the separation of Native American ancestors. A more parsimonious explanation for the present-day distribution of the haplotype is that Native Americans retain the ancestral state of Paleolithic Eurasians. Another haplotype at the locus may reflect a secondary selection signal, although its functional impact is unknown.

## Introduction

Long-chain polyunsaturated fatty acids (LC-PUFA) are essential for many aspects of cellular and organismal function (Marszalek and Lodish 2005; Darios and Davletov 2006). While they can be obtained from dietary sources, they can also be synthesized from short-chain PUFA (SC-PUFA) through the ω-3 and ω-6 biosynthesis pathways. Some of the steps in these pathways are catalyzed by the fatty acid desaturase genes *FADS1* and *FADS2* (Nakamura and Nara 2004) which are located close to each other on human chromosome 11. This locus (which we refer to as the *FADS* locus) has been targeted by selection multiple times in human evolution (Ameur, et al. 2012; Mathias, et al. 2012; Mathieson, et al. 2015; Buckley, et al. 2017; Ye, et al. 2017; Mathieson and Mathieson 2018). There are two LD blocks at the locus, but most studies have focused on the two major haplotypes at LD block 1 (Ameur, et al. 2012), which we refer to as the *ancestral* (A) and *derived* (D) haplotypes. LD block 1 is also where the strongest genome-wide association study signals for lipid levels are detected in European ancestry populations (Teslovich, et al. 2010). Haplotype D appears to have been under selection–likely preceding the out-of-Africa bottleneck–in Africa, and is virtually fixed in present-day African populations (Ameur, et al. 2012; Mathias, et al. 2012). Given this, it is surprising that early Eurasian populations appear to have largely carried the ancestral haplotype, suggesting selection for the ancestral haplotype at some time after the split of present-day African and non-African lineages (Ye, et al. 2017; Mathieson and Mathieson 2018). By the Mesolithic–around 10,000 years before present (BP)–the ancestral haplotype was fixed in Europe (Mathieson, et al. 2015). The derived haplotype was reintroduced to Europe in the Neolithic (beginning around 8400 BP) by the migration of Early Farming populations from Anatolia (Mathieson, et al. 2015), experienced strong positive selection during the Bronze Age (Buckley, et al. 2017; Mathieson and Mathieson 2018), and is now at a frequency of around 60%. The trajectory of the derived haplotype in East Asian populations is less clear, but the observations that the 40,000 year old Tianyuan individual is homozygous for the ancestral haplotype and that the locus has a strong signal of selection in East Asian populations (1000 Genomes Project Consortium 2015), suggest that a very similar process might have occurred.

### The ancestral haplotype was common in Upper Paleolithic Eurasia

We examined ancient DNA from 16 individuals from Early Upper Paleolithic Eurasia (Fu, et al. 2014; Raghavan, et al. 2014; Fu, et al. 2016; Sikora, et al. 2017; Yang, et al. 2017; Sikora, et al. 2019). Of these individuals’ 32 haplotypes, 4 are derived and 28 are ancestral (3 vs 23 supported by more than 6 reads; Figure 1e). This confirms that the derived LD block 1 haplotype was uncommon, though not completely absent, throughout Upper Paleolithic Eurasia. With a sufficiently intense bottleneck, genetic drift could fix the ancestral haplotype in the ancestors of Native Americans even if it was not under selection (Harris, et al. 2019).

**Figure 1:**
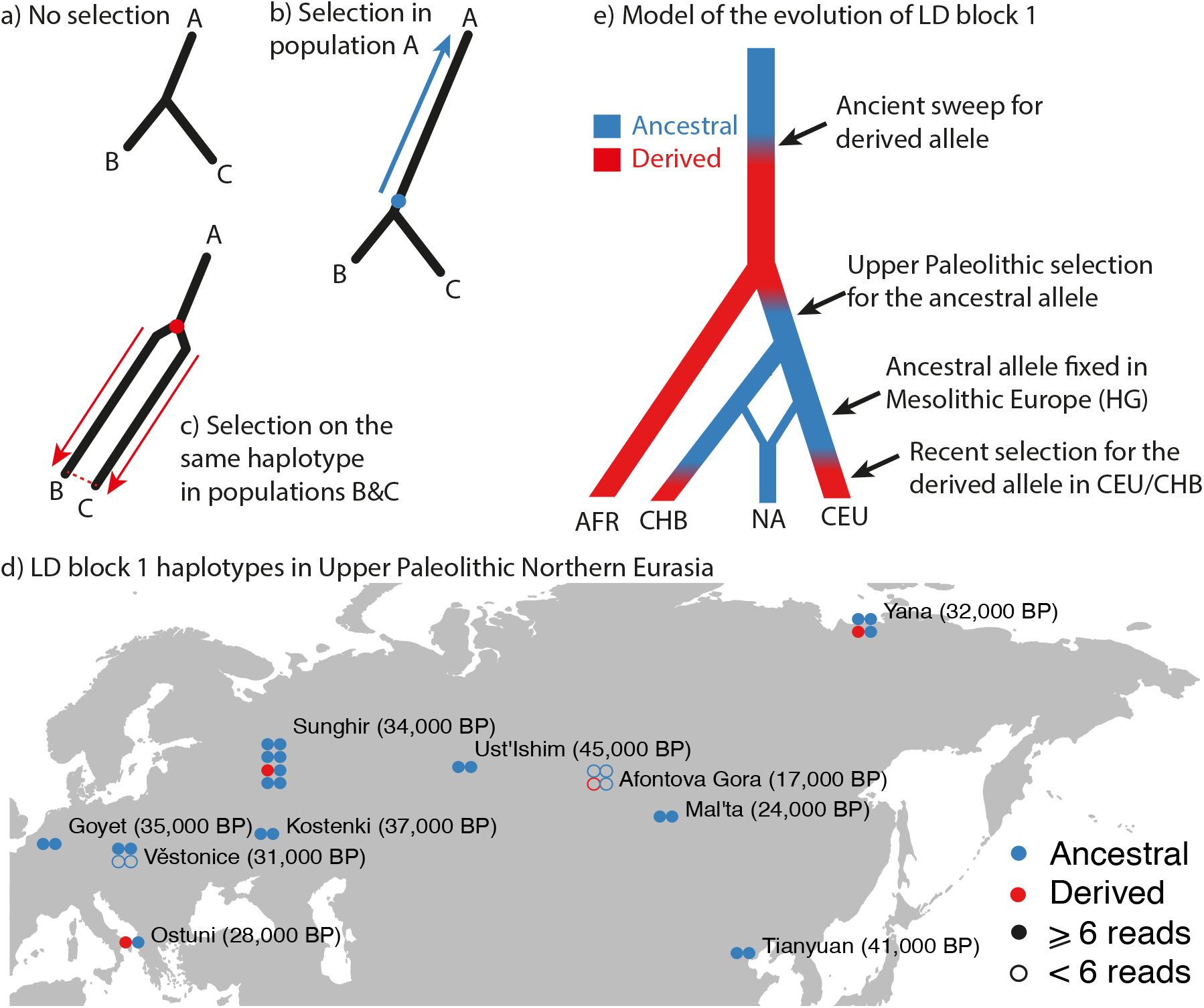
**a**) The PBS compares genetic differentiation (branch length) between three populations. **b**) If a new mutation (blue dot) is under selection in one population (A), the branch length leading to A will be longer–a signal of selection. **c**). If a haplotype that exists in the ancestral population (red dot) is under selection in both populations B and C then, because B and C look very similar, the PBS misattributes the long branch length to A, instead of to B and C. **d**) LD block 1 haplotypes in Upper Paleolithic Eurasia (Fu, et al. 2014; Raghavan, et al. 2014; Fu, et al. 2016; Sikora, et al. 2017; Yang, et al. 2017; Sikora, et al. 2019). Ancestral and derived haplotype defined as haplotypes A and C (Mathieson and Mathieson 2018) **e**) Model for the evolution of present-day African (AFR), East Asian (CHB), Native American (NA) and European (CEU) population showing where derived (red) and ancestral (blue) haplotypes are common.

### Selection scans at the locus are confounded by parallel adaptation

Excluding the effects of recent admixture, present-day Native American, Inuit, and Siberian populations are fixed for the ancestral haplotype (Ameur, et al. 2012; Fumagalli, et al. 2015; Harris, et al. 2019). Fumagalli, et al. (2015) found a strong signal of selection at the locus in the Greenlandic Inuit population, which they interpret as an adaption to a PUFA-rich Arctic diet at least 20,000 years ago in the common ancestors of present-day Inuit and Siberian populations. Subsequent studies detected a very similar signal in Native American populations (Amorim, et al. 2017; Hlusko, et al. 2018; Harris, et al. 2019). Some studies interpret this as evidence for selection for the ancestral haplotype relatively early on the Native American lineage, although it could also represent a shared signal from the population ancestral to both Native Americans and Inuit. Both these signals extend across LD blocks 1 (tagging the ancestral haplotype) and 2.

These studies used the population branch statistic (PBS) (Yi, et al. 2010) to compare Native American (NA), European (EUR) and East Asian (EAS) populations. The PBS compares genetic differentiation between three populations and identifies which, if any, of the three branches has excess differentiation that indicates selection. We write PBS(A, (B, C)) to denote the statistic that is testing for excess differentiation on the A branch, relative to B and C. However, the PBS makes the implicit assumption that each of the three branches is independent. If there is parallel selection on the same haplotype in two population, say B and C, then B and C will have low F_ST_, but each will have high F_ST_ relative to A. Thus, the PBS will misattribute selection to branch A (Fig. 1a-c). Given low frequency of the derived allele in Upper Paleolithic Eurasia (Fig. 1d) and parallel selection in EUR and EAS populations (Fig. 1e), we expect that PBS(NA, (EUR, EAS)) would give a spurious signal of selection in the Native American population.

We tested this by computing the PBS using Native American, European, and Mesolithic European populations. We used the PEL (Peruvians from Lima) 1000 Genomes population to represent Native Americans (1000 Genomes Project Consortium 2015). For each PEL individual, we restricted to regions of homozygous Native American ancestry (Martin, et al. 2017). We used CEU (Northern and Western European ancestry) to represent present-day Europeans, CHB (Chinese from Beijing) to represent East Asians and 118 ancient European hunter-gatherers (HG) to represent Mesolithic Europe (Mathieson and Mathieson 2018). We replicate the elevated PBS(PEL, (CEU, CHB)) statistic at the *FADS* locus (Fig.2 left column; upper 0.0002 quantile), but find that it largely disappears if we replace CHB with HG (Fig. 2 right column; upper 0.016 quantile). Two of the LD block 2 SNPs originally identified by Fumagalli, et al. (2015)–rs74771917 and rs7115739–still have extreme PBS values, but the PBS is no longer elevated across the locus (Fig.2). Conversely, both PBS(CEU, (PEL, HG)) and PBS(CHB, (PEL, HG)) do show elevated values (upper 0.00004 and 0.0006 quantiles). These results are consistent with the model of parallel adaptation in CEU and CHB shown in Fig. 1d.

**Figure 2:**
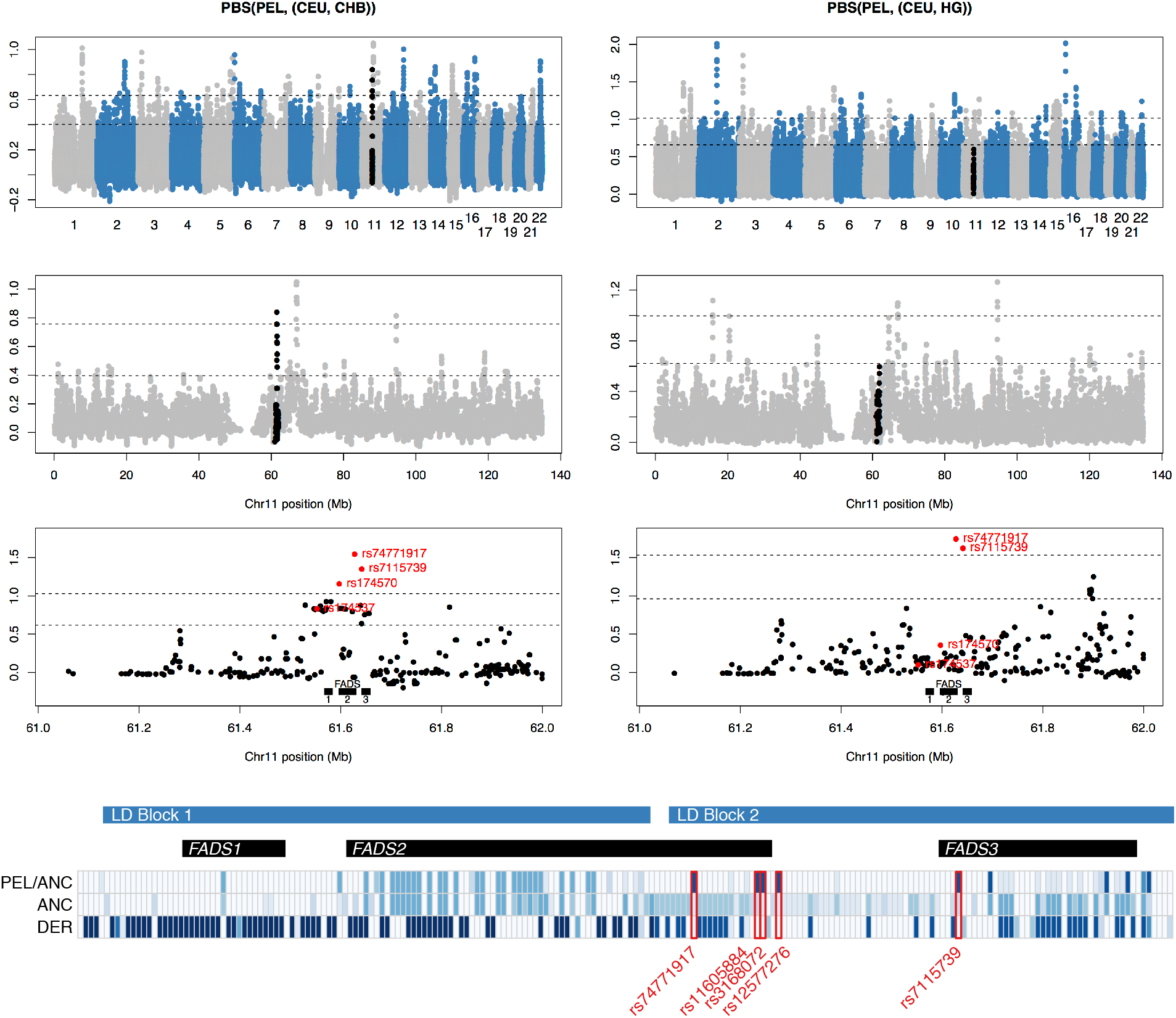
PBS on the Native American branch. Left column; PBS(PEL,(CEU,CHB)). Right column, PBS(PEL,(CEU,HG)). Upper row; genome-wide PBS in overlapping 20-SNP windows, each shifted by 5 SNPs. Black points indicate the region Chr11:61-62Mb (hg19) that contains the *FADS* locus. Second row; Chromosome 11 PBS in overlapping 20-SNP windows. Third row; per-SNP PBS in the region Chr11:61-62M. Horizontal lines indicate upper 0.01 and 0.001 genome-wide PBS quantiles. In the lower row, red labeled points indicate SNPs previously identified as targets of selection (Fumagalli, et al. 2015; Amorim, et al. 2017). Top three rows restricted to 903,961 autosomal SNPs present on the 1240k capture array with a minor allele frequency of at least 5% in at least one of the four populations. Lower row: Allele frequencies for all SNPs at >1% frequency in at least one population in CEU and CHB individuals carrying the derived haplotype (DER), the ancestral haplotype (ANC) and for PEL individuals carrying the ancestral haplotype (PEL/ANC). Color indicates the frequency of the variant that is less common on the ancestral haplotype. Highlighted in red are the five LD block 2 SNPs that have >50% difference in frequency between ANC and PEL/ANC, and at most 10% frequency in DER.

### A potential secondary signal of selection

While the Native American PBS signal in LD block 1 (containing rs174570, rs174556 and rs174537) disappears when Mesolithic hunter-gatherers are used as an outgroup, part of the signal (at rs74771917 and rs7115739) that is restricted to LD block 2 remains (Fig. 2). Using sequence data from the 1000 Genomes project we identified three additional SNPs that are highly differentiated between present-day Native American and Eurasian ancestral allele carriers (Figure 4). Among individuals who carry the ancestral haplotype at LD block 1, this LD block 2 haplotype has a frequency of around 100% in the Greenland Inuit (Fumagalli, et al. 2015) 82% in PEL, 34% in CHB, 9% in HG and 4% in CEU. The Anzick individual (Rasmussen, et al. 2014) carries two copies, while the 40,000-year old East Asian Tianyuan individual (Yang, et al. 2017) carries one. It therefore remains possible that the high frequency of this haplotype represents a secondary signal of selection in the common ancestor of Inuit and Native Americans. On the other hand, this region is not a genome-wide outlier in the standard window-based PBS analysis and the frequency of this haplotype may have been driven by linked selection on LD block 1. Further, the LD block 2 haplotype has not been shown to affect expression of any of the *FADS* genes or any other phenotype, independent of the LD block 1 haplotype. Within-population, the two blocks are highly correlated, so it would be necessary to perform conditional analysis at the locus in East Asian populations to identify any independent effect of the LD block 2 haplotype.

## Conclusion

The Native American specific signal of selection at the FADS locus is largely an artefact driven by parallel selection on the European and East Asian lineages. The ancestral haplotype at LD block 1 may have been selected in Upper Paleolithic Eurasian populations but this likely took place around or before the split of East and West Eurasian populations 40-60,000 years ago and certainly before the Native American and Siberian lineages split. There remains some evidence of a secondary signal of selection in LD block 2 but this is shared by Inuit and Siberians and not specific to Native Americans. The complex history of selection at this locus likely confounds selection scans in other populations as well. Finally, this analysis demonstrates the ability of direct evidence from ancient DNA to resolve complex evolutionary histories that may not be identifiable using present-day data.

## Acknowledgments

I.M. was supported by a Research Fellowship from the Alfred P. Sloan foundation [FG-2018-10647], a New Investigator Research Grant from the Charles E. Kaufman Foundation [KA2018-98559], and NIGMS award number [R35GM133708]. The content is solely the responsibility of the author and does not necessarily represent the official views of the National Institutes of Health. We thank Shai Carmi, Matteo Fumagalli, Rasmus Nielsen, Fernando Racimo and Pontus Skoglund for helpful comments on earlier drafts.

---

**Supplementary Figure 1:**
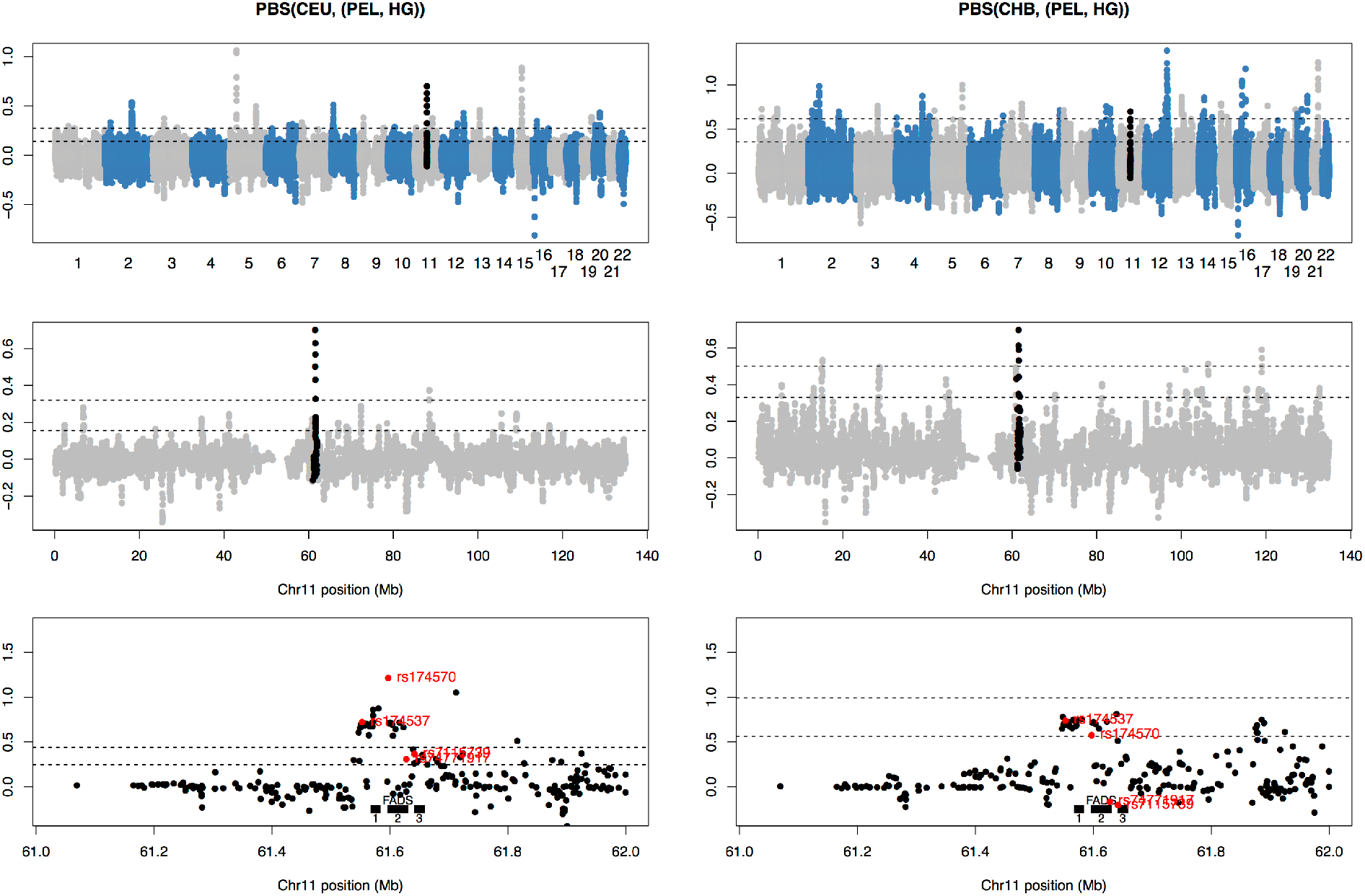
Population branch statistics around the *FADS* locus on European and East Asian branches. Left column; PBS(CEU,(PEL,HG)). Right column, PBS(CHB,(PEL,HG)). Upper row; genome-wide PBS in overlapping 20-SNP windows, each shifted by 5 SNPs. Black points indicate the region Chr11:61-62Mb (hg19) that contains the *FADS* locus. Middle row; Chromosome 11 PBS in overlapping 20-SNP windows. Lower row; per-SNP PBS in the region Chr11:61-62M. In each plot, horizontal lines indicate upper 0.01 and 0.001 genome-wide PBS quantiles. In the lower row, the location of *FADS1, 2* and 3 is indicted with black bars, while red labeled points indicate SNPs previously identified as targets of selection (Fumagalli, et al. 2015; Amorim, et al. 2017). Restricted to 903,961 SNPs present on the 1240k capture array with a minor allele frequency of at least 5% in at least one of the four populations.

**Supplementary Figure 2:**
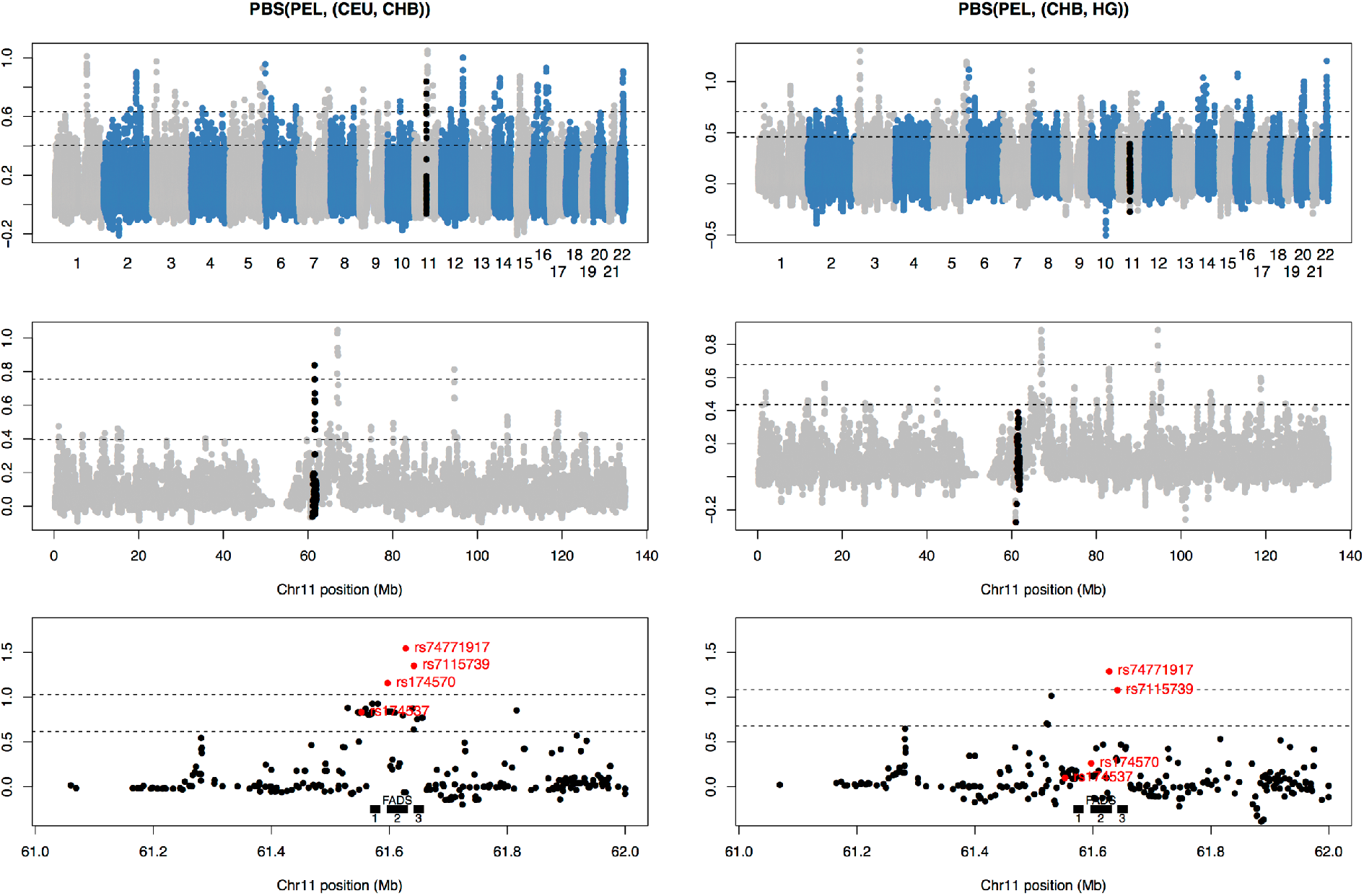
As Figure 2, but with PBS(PEL,(CHB,HG)) in the right-hand column, instead of PBS(PEL, (CEU,HG)

**Supplementary Table 1:** Reads supporting the reference (derived) and alternative (ancestral) allele (der,anc) at 11 SNPs used to define derived haplotype C (Mathieson and Mathieson 2018).

| CHR | POS | ID | REF | ALT | GoyetQ116 | Vestonice 13 | Vestonice 16 | Ostuni 1 | Kostenki 14 | Sungir I | Sungir II | Sungir III | Sungir IV | Afontova Gora 2 | Afontova Gora 3 | Ust'Ishim | MA1 | Tianyuan | Yana | Yana 2 |
| --- | --- | --- | --- | --- | --- | --- | --- | --- | --- | --- | --- | --- | --- | --- | --- | --- | --- | --- | --- | --- |
| 11 | 61551927 | rs174536 | A | C | 0,1 | 0,2 | 0,1 | 1,1 | 0,12 | 0,1 | 0,4 | 3,5 | 0,1 | 0,0 | 0,0 | 0,29 | 0,1 | 0,0 | 11,7 | 0,8 |
| 11 | 61552680 | rs174537 | G | T | 0,3 | 0,0 | 0,0 | 0,0 | 0,16 | 0,1 | 0,6 | 3,8 | 0,2 | 1,0 | 0,0 | 0,19 | 0,1 | 0,5 | 11,10 | 0,8 |
| 11 | 61557826 | rs102274 | T | C | 0,6 | 0,3 | 0,10 | 1,2 | 0,93 | 0,1 | 0,7 | 10,7 | 0,7 | 0,0 | 0,0 | 0,20 | 0,1 | 0,11 | 10,20 | 0,4 |
| 11 | 61569306 | rs174545 | C | G | 0,0 | 0,0 | 0,0 | 0,0 | 0,0 | 0,2 | 0,5 | 6,6 | 0,3 | 0,0 | 0,0 | 0,39 | 0,0 | 0,0 | 14,20 | 0,8 |
| 11 | 61569830 | rs174546 | C | T | 0,6 | 0,0 | 0,17 | 0,0 | 0,75 | 0,0 | 0,4 | 7,8 | 0,7 | 0,1 | 0,2 | 0,38 | 0,4 | 0,0 | 10,14 | 0,8 |
| 11 | 61570783 | rs174547 | T | C | 0,0 | 0,0 | 0,5 | 0,0 | 1,19 | 0,0 | 0,9 | 9,3 | 0,4 | 0,0 | 0,0 | 0,33 | 0,0 | 0,7 | 13,15 | 1,7 |
| 11 | 61571478 | rs174550 | T | C | 0,0 | 0,0 | 0,10 | 0,0 | 0,67 | 0,5 | 0,5 | 5,7 | 0,5 | 0,1 | 0,1 | 0,25 | 0,1 | 0,9 | 11,17 | 0,6 |
| 11 | 61575158 | rs174553 | A | G | 0,0 | 0,0 | 0,0 | 0,0 | 0,0 | 0,0 | 0,8 | 3,7 | 0,3 | 1,0 | 0,0 | 0,30 | 0,0 | 0,0 | 9,17 | 0,4 |
| 11 | 61579463 | rs174554 | A | G | 0,0 | 0,0 | 0,3 | 1,0 | 1,20 | 0,0 | 0,2 | 4,4 | 0,4 | 0,0 | 0,0 | 5,20 | 0,0 | 0,0 | 14,16 | 1,6 |
| 11 | 61585144 | rs174562 | A | G | 0,0 | 0,0 | 0,0 | 0,0 | 0,0 | 0,1 | 0,4 | 10,9 | 0,4 | 0,0 | 0,0 | 1,46 | 0,0 | 0,0 | 21,18 | 0,3 |
| 11 | 61588305 | rs174564 | A | G | 0,0 | 0,0 | 2,0 | 0,0 | 7,9 | 0,0 | 0,1 | 3,1 | 0,5 | 0,0 | 0,0 | 15,15 | 0,0 | 0,0 | 7,1 | 0,2 |
| Ancestral (A) or Dervied (D) haplotype |  |  |  |  | AA | AA? | AA | AD | AA | AA | AA | AD | AA | AD? | AA? | AA | AA | AA | AD | AA |

## References

1000 Genomes Project Consortium. 2015. A global reference for human genetic variation. Nature 526:68–74.

Ameur A, Enroth S, Johansson A, Zaboli G, Igl W, Johansson AC, Rivas MA, Daly MJ, Schmitz G, Hicks AA, et al. 2012. Genetic adaptation of fatty-acid metabolism: a human-specific haplotype increasing the biosynthesis of long-chain omega-3 and omega-6 fatty acids. Am J Hum Genet 90:809–820.

Amorim CE, Nunes K, Meyer D, Comas D, Bortolini MC, Salzano FM, Hunemeier T. 2017. Genetic signature of natural selection in first Americans. Proc Natl Acad Sci U S A 114:2195–2199.

Buckley MT, Racimo F, Allentoft ME, Jensen MK, Jonsson A, Huang H, Hormozdiari F, Sikora M, Marnetto D, Eskin E, et al. 2017. Selection in Europeans on Fatty Acid Desaturases Associated with Dietary Changes. Mol Biol Evol 34:1307–1318.

Darios F, Davletov B. 2006. Omega-3 and omega-6 fatty acids stimulate cell membrane expansion by acting on syntaxin 3. Nature 440:813–817.

Fu Q, Li H, Moorjani P, Jay F, Slepchenko SM, Bondarev AA, Johnson PL, Aximu-Petri A, Prufer K, de Filippo C, et al. 2014. Genome sequence of a 45,000-year-old modern human from western Siberia. Nature 514:445–449.

Fu Q, Posth C, Hajdinjak M, Petr M, Mallick S, Fernandes D, Furtwängler A, Haak W, Meyer M, Mittnik A, et al. 2016. The genetic history of Ice Age Europe. Nature 534:200–205.

Fumagalli M, Moltke I, Grarup N, Racimo F, Bjerregaard P, Jorgensen ME, Korneliussen TS, Gerbault P, Skotte L, Linneberg A, et al. 2015. Greenlandic Inuit show genetic signatures of diet and climate adaptation. Science 349:1343–1347.

Harris DN, Ruczinski I, Yanek LR, Becker LC, Becker DM, Guio H, Cui T, Chilton FH, Mathias RA, O’Connor T. 2019. Evolution of Hominin Polyunsaturated Fatty Acid Metabolism: From Africa to the New World. Genome Biology and Evolution 11:1417–1430.

Hlusko LJ, Carlson JP, Chaplin G, Elias SA, Hoffecker JF, Huffman M, Jablonski NG, Monson TA, O’Rourke DH, Pilloud MA, et al. 2018. Environmental selection during the last ice age on the mother-to-infant transmission of vitamin D and fatty acids through breast milk. Proc Natl Acad Sci U S A.

Marszalek JR, Lodish HF. 2005. Docosahexaenoic acid, fatty acid-interacting proteins, and neuronal function: breastmilk and fish are good for you. Annu Rev Cell Dev Biol 21:633–657.

Martin AR, Gignoux CR, Walters RK, Wojcik GL, Neale BM, Gravel S, Daly MJ, Bustamante CD, Kenny EE. 2017. Human Demographic History Impacts Genetic Risk Prediction across Diverse Populations. Am J Hum Genet 100:635–649.

Mathias RA, Fu W, Akey JM, Ainsworth HC, Torgerson DG, Ruczinski I, Sergeant S, Barnes KC, Chilton FH. 2012. Adaptive evolution of the FADS gene cluster within Africa. PLoS One 7:e44926.

Mathieson I, Lazaridis I, Rohland N, Mallick S, Patterson N, Roodenberg SA, Harney E, Stewardson K, Fernandes D, Novak M, et al. 2015. Genome-wide patterns of selection in 230 ancient Eurasians. Nature 528:499–503.

Mathieson S, Mathieson I. 2018. FADS1 and the Timing of Human Adaptation to Agriculture. Mol Biol Evol 35:2957–2970.

Nakamura MT, Nara TY. 2004. Structure, function, and dietary regulation of delta6, delta5, and delta9 desaturases. Annu Rev Nutr 24:345–376.

Raghavan M, Skoglund P, Graf KE, Metspalu M, Albrechtsen A, Moltke I, Rasmussen S, Stafford Jr TW, Orlando L, Metspalu E, et al. 2014. Upper Palaeolithic Siberian genome reveals dual ancestry of Native Americans. Nature 505:87–91.

Rasmussen M, Anzick SL, Waters MR, Skoglund P, DeGiorgio M, Stafford Jr TW, Rasmussen S, Moltke I, Albrechtsen A, Doyle SM, et al. 2014. The genome of a Late Pleistocene human from a Clovis burial site in western Montana. Nature 506:225–229.

Sikora M, Pitulko VV, Sousa VC, Allentoft ME, Vinner L, Rasmussen S, Margaryan A, de Barros Damgaard P, de la Fuente C, Renaud G, et al. 2019. The population history of northeastern Siberia since the Pleistocene. Nature 570:182–188.

Sikora M, Seguin-Orlando A, Sousa VC, Albrechtsen A, Korneliussen T, Ko A, Rasmussen S, Dupanloup I, Nigst PR, Bosch MD, et al. 2017. Ancient genomes show social and reproductive behavior of early Upper Paleolithic foragers. Science 358:659–662.

Teslovich TM, Musunuru K, Smith AV, Edmondson AC, Stylianou IM, Koseki M, Pirruccello JP, Ripatti S, Chasman DI, Willer CJ, et al. 2010. Biological, clinical and population relevance of 95 loci for blood lipids. Nature 466:707–713.

Yang MA, Gao X, Theunert C, Tong H, Aximu-Petri A, Nickel B, Slatkin M, Meyer M, Pääbo S, Kelso J, et al. 2017. 40,000-Year-Old Individual from Asia Provides Insight into Early Population Structure in Eurasia. Current Biology 27:3202–3208.e3209.

Ye K, Gao F, Wang D, Bar-Yosef O, Keinan A. 2017. Dietary adaptation of FADS genes in Europe varied across time and geography. Nat Ecol Evol 1:167.

Yi X, Liang Y, Huerta-Sanchez E, Jin X, Cuo ZX, Pool JE, Xu X, Jiang H, Vinckenbosch N, Korneliussen TS, et al. 2010. Sequencing of 50 human exomes reveals adaptation to high altitude. Science 329:75–78.

